# Mobile-CRISPRi as a powerful tool for modulating *Vibrio* gene expression

**DOI:** 10.1101/2024.01.17.575898

**Authors:** Logan Geyman, Madeline Tanner, Natalia Rosario-Melendez, Jason Peters, Mark J. Mandel, Julia C. van Kessel

## Abstract

CRISPRi (Clustered Regularly Interspaced Palindromic Repeats interference) is a gene knockdown method that uses a deactivated Cas9 protein (dCas9) that binds a specific gene target locus dictated by an encoded guide RNA (sgRNA) to block transcription. Mobile-CRISPRi is a suite of modular vectors that enable CRISPRi knockdowns in diverse bacteria by integrating IPTG-inducible *dcas9* and *sgRNA* genes into the genome using Tn*7* transposition. Here, we show that the Mobile-CRISPRi system functions robustly and specifically in multiple *Vibrio* species: *Vibrio cholerae, Vibrio fischeri, Vibrio vulnificus, Vibrio parahaemolyticus*, and *Vibrio campbellii*. We demonstrate efficacy by targeting both essential and non-essential genes that function to produce defined, measurable phenotypes: bioluminescence, quorum sensing, cell division, and growth arrest. We anticipate that Mobile-CRISPRi will be used in *Vibrio* species to systematically probe gene function and essentiality in various behaviors and native environments.

## Introduction

The rise of Clustered Regularly Interspaced Palindromic Repeats (CRISPR) as a genetic tool has empowered researchers to make precise, controlled changes to the genomes of model and non-model organisms [1]. The development of a CRISPR based interference tool (CRISPRi) has allowed researchers to modulate gene expression [2-5]. CRISPRi works by targeting an enzymatically inactivated CRISPR associated protein 9 (dCas9) to a gene of interest via a complementary single guide RNA (sgRNA); the mutation in *dcas9* prevents double-stranded cleavage of the DNA but the dCas9 protein remains tightly associated with its DNA target [6]. This protein complex prevents transcription by inhibiting the initiation or progression of RNA polymerase at the target locus [6]. The gene of interest is dictated via introduction of a specific modifiable sequence in the sgRNA called a spacer that is complementary to the sense strand of the target (protospacer). In this form of Mobile-CRISPRi, the suite of CRISPRi machinery includes type-IIA *dcas9*, an sgRNA cloning region, and inducible P_*lacO-1*_ promoters driving expression of both, and the entire suite has been grafted onto a Tn*7* transposon plasmid capable of integration into a conserved chromosomal attachment site in a broad range of target hosts (Fig. 1A) [2, 3, 7-9]. Thus, the entire system is permanently integrated into the bacterial chromosome, eliminating the need for antibiotic selection during propagation, which is important for use of these strains in host infection assays [9]. The Mobile-CRISPRi suite is highly modular, allowing researchers to customize the promoters that control expression of the *dcas9* and sgRNA. Additionally, there are alternate forms of the system for organisms that are incompatible with Tn7 transposition [2, 3]. The initial Mobile-CRISPRi study demonstrated gene knockdown in cultured *Vibrio casei* cells, but not other *Vibrio* species [3]. Those data, coupled with the potential for wide ranging applications including studies in animal hosts, make Mobile-CRISPRi an especially valuable resource to develop for use across *Vibronaceae*.

**Fig. 1.**
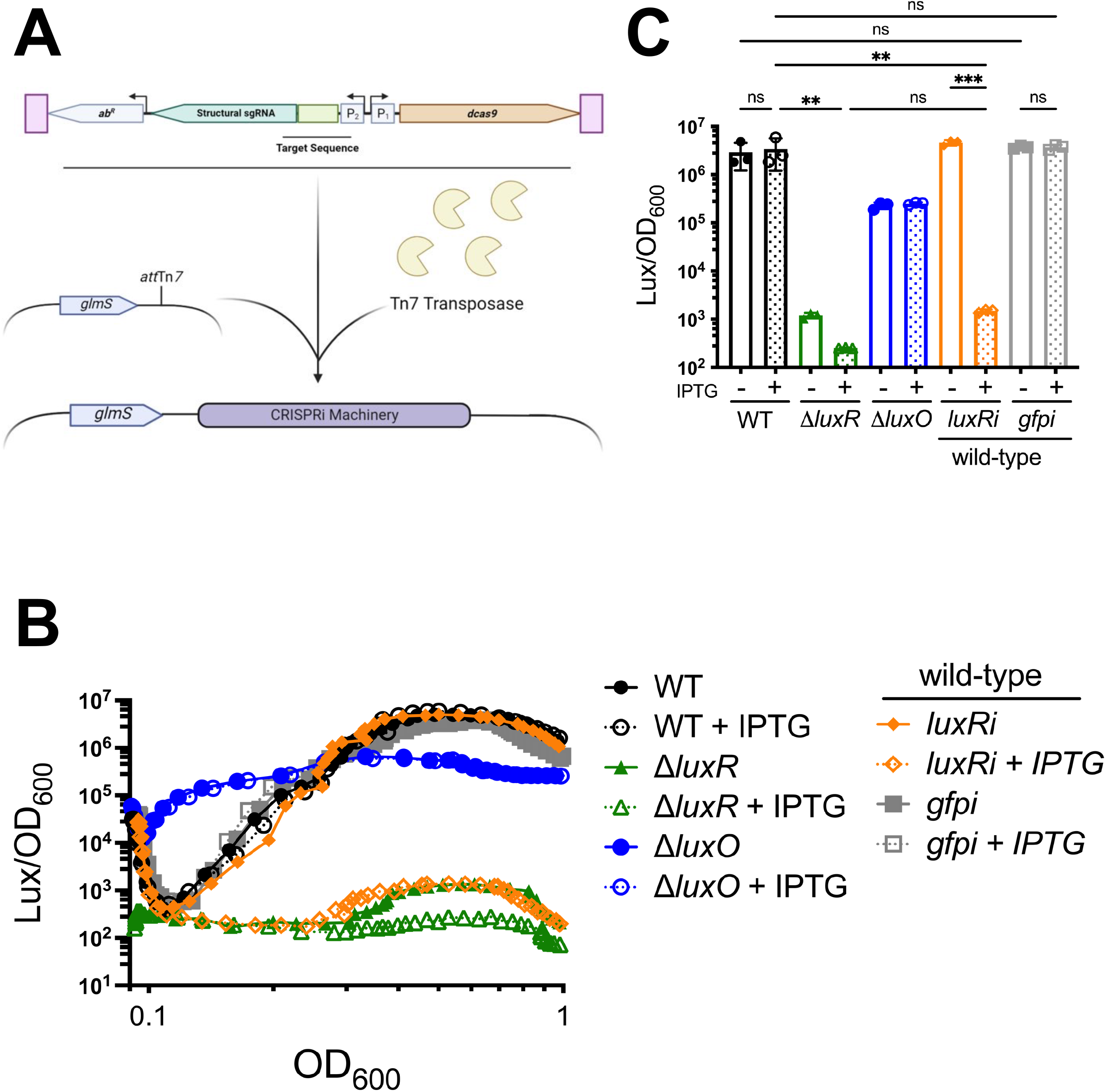
Mobile CRISPRi targets *luxR* in *V. campbellii*. (A) Model depicting Tn*7* insertion of Mobile-CRISPRi with a variable targeting sequence. (B) Bioluminescence production (Lux/OD_600_) throughout a growth curve or (C) endpoint production at 15 hours is shown for *V. campbellii* strains BB120 (WT), ti*luxR*, ti*luxO*, or wild-type containing the CRISPRi plasmid targeting either *luxR* (pLG002) or *gfp* (pLG027) in the presence or absence of 100 μM IPTG. (B) A representative light curve of three biological replicates is shown. (C) Error bars represent standard deviation, and the gene diagram shows the relative location of the sgRNA used. Statistical analysis: normally distributed data (Shapiro-Wilk test) were analyzed by one-way analysis of variance (ANOVA; *n*=3; Tukey’s multiple comparisons test; **, *p*=0.01; ***, *p*=0.001; ns=not significant). Select comparisons are shown because not all could be graphed simultaneously.

The aim of this work was to determine the efficacy of Mobile-CRISPRi in *Vibrio* bacteria. This diverse group includes both symbiotic (*Vibrio fischeri*) and pathogenic species (*Vibrio cholerae, Vibrio vulnificus, Vibrio parahaemolyticus, Vibrio campbellii, Vibrio coralliilyticus*), which are relevant to human health, biotechnology, aquaculture, and marine ecology [10-12]. *Vibrio* species are the model organism subjects of research in a plethora of research fields: natural transformation, type III secretion, type VI secretion, bacterial motility and adherence, biofilm formation, quorum sensing, bacteriophage-host evolution, chromosome and plasmid replication, host-mutualist interactions, host-pathogen interactions, and more [13-18]. Here, we assessed the efficacy of Mobile-CRISPRi by targeting several systems for gene expression knockdown. First, we targeted the gene encoding the master quorum sensing transcriptional regulator LuxR in *V. campbellii*, which activates bioluminescence at high cell density [10, 16, 17, 19]. We used this system to optimize induction of the system and characterize the effect of sgRNA location within an open reading frame. We also determined the utility of CRISPRi in studying essential genes, including the gene encoding β-subunit of RNA polymerase (*rpoB*) and the cell division gene *ftsZ* [20, 21]. Finally, we show that Mobile-CRISPRi is effective in multiple *Vibrio* species that are frequently studied as model organisms, indicating the broad applicability of this system for genetic research.

## Results

### *Mobile-CRISPRi functions to specifically target genes in* V. campbellii

To assess the function of Mobile-CRISPRi in *Vibrio*, we began our studies using an easily quantifiable phenotype: bioluminescence. In *V. campbellii*, the master transcriptional regulator LuxR is required for the transcriptional activation of the bioluminescence gene operon *luxCDABE* [10, 16, 17, 22]. Therefore, we chose *luxR* as the first target gene to assess the effectiveness of Mobile-CRISPRi and designed a guide RNA spacer to target *luxR* near the 5’ end of the transcript (designated *luxRi*) [6, 23]. To assess non-specific knockdown effects on gene expression, we also designed a guide RNA to target the *gfp* gene (designated *gfpi*) in cells with and without the *gfp* gene. These Mobile-CRISPRi plasmids were conjugated into *V. campbellii* BB120, and these strains were assayed for bioluminescence in the presence or absence of IPTG inducer. As controls for bioluminescence production, we also assayed a constitutively bioluminescent strain (Δ*luxO*) and a non-bioluminescent strain (Δ*luxR*) alongside the CRISPRi strains. We observed that addition of IPTG to the *luxRi* strain resulted in loss of bioluminescence production throughout the growth curve, displaying the same phenotype as the Δ*luxR* strain (Fig. 1B, 1C). The CRISPRi effect on *luxR* became saturating at 10 µM of IPTG (Fig. S1A). Induction of the *luxRi* CRISPRi genes in the *luxRi* strain did not alter growth (Fig. S2). Further, induction of the *gfpi* targeting sgRNA did not alter the production of bioluminescence in the presence or absence of IPTG, indicating that the sgRNA effects are specific (Fig. 1B, 1C). We observe that there is no difference in bioluminescence levels in uninduced wild-type and uninduced *luxRi* (Fig. 1C), suggesting there is little to no leakiness in the system within this context. From these data, we conclude that Mobile-CRISPRi specifically targets gene loci in *V. campbellii* without any observable growth or off-target effects of active dCas9-guide RNA complexes.

### Guide RNA location impacts knockdown efficacy

Previous studies have demonstrated the potential utility in altering CRISPRi by targeting the guide RNA to different sections of a gene’s open reading frame [6, 23]. To assess this modification in *V. campbellii* BB120, we developed *luxRi* strains targeting different sections of the open reading frame (Fig. 2A). The same controls as the previous assay were used. We observed that the repression conveyed by CRISPRi steadily decreased as the guide RNA target moved from the 5’ end toward the 3’ in the open reading frame. The observation was recapitulated in wild-type strains targeting *luxC (luxCi)* (Fig. 2B). These data suggest that achieving high efficacy gene knockdown in *Vibrio* species necessitates targeting towards the 5’ end of the open reading frame.

**Fig. 2.**
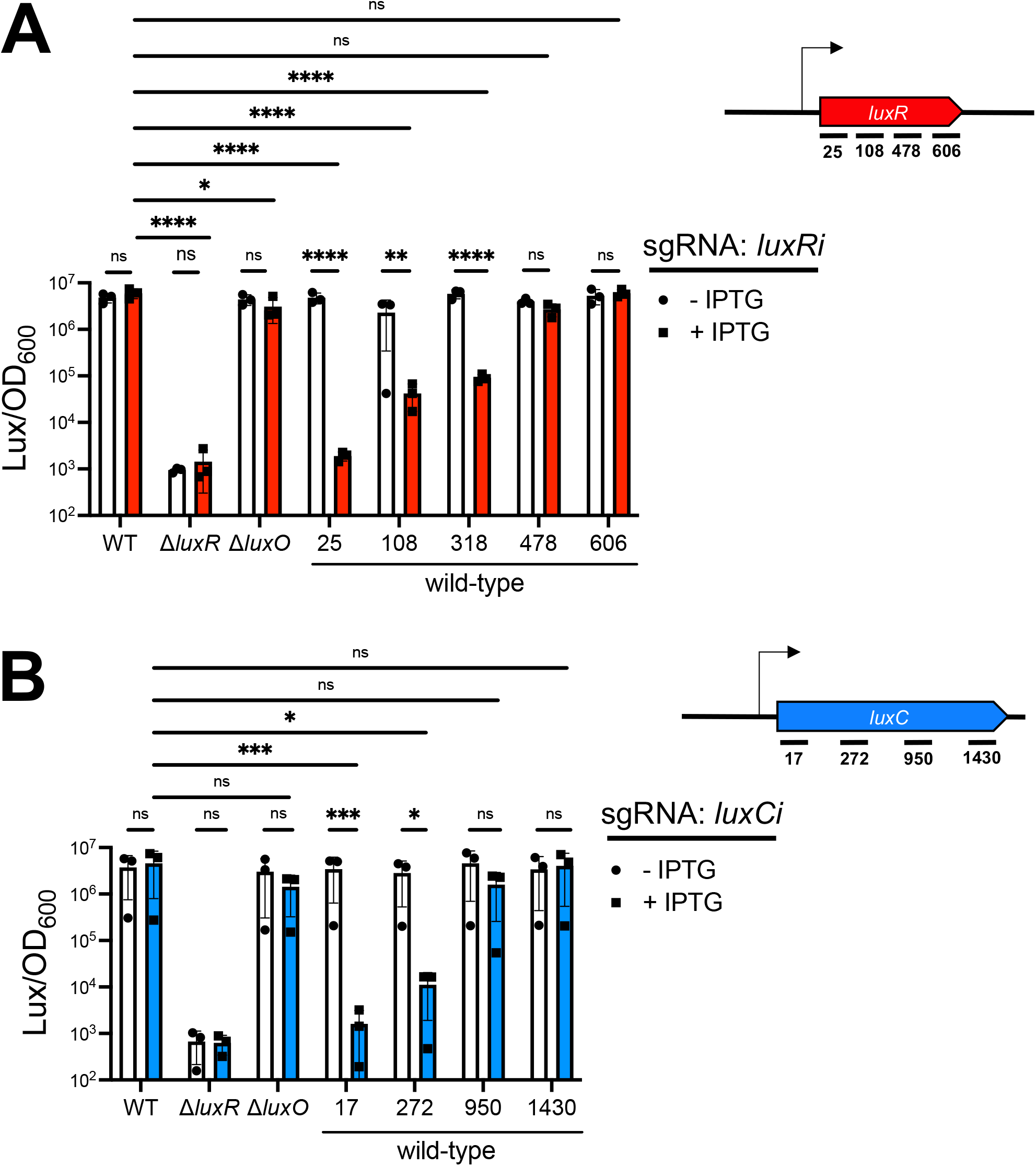
Effects of guide RNA location. Bioluminescence endpoint production (Lux/OD_600_) at 15 hours is shown for *V. campbellii* strains BB120 (WT), t<*luxR*, t<*luxO*, or wild-type containing a CRISPRi plasmid with guide RNA targeting *luxR* (A) or *luxC* (B) in the presence or absence of 100 μM IPTG. The location of the guide RNA target from the beginning of the open reading frame (ORF) is indicated in base-pairs. Next to each panel is a gene diagram depicting the relative location of each sgRNA along the ORF denoted by the same base-pair number. Statistical analysis: normally distributed data (D’Agostino and Pearson test) were analyzed by two-way analysis of variance of log-transformed data (*n*=3; Tukey’s multiple comparisons test; *, *p*=0.05; ***, *p*=0.001; ****, *p*=0.0001; ns=not significant). Select comparisons are shown because not all could be graphed simultaneously.

### Mobile-CRISPRi targets the essential genes rpoB and ftsZ

A significant value in applying CRISPRi compared to a gene inactivation approach (e.g., deletion) is the ability examine essential genes. To assess the utility of the CRISPRi system in temporal knockdown of essential genes, we targeted the gene encoding the β subunit of RNA polymerase (*rpoB)* [20, 21]. Upon induction of the *V. campbellii rpoBi* strain with IPTG, we observed a severely delayed lag phase compared to the wild-type strain and the control *rpoBi* strain without inducer (Fig. 3A). Although growth eventually recovered for the *rpoBi* strain, we hypothesize that this growth is likely due to suppressor mutants as has been demonstrated previously [24].

**Fig. 3.**
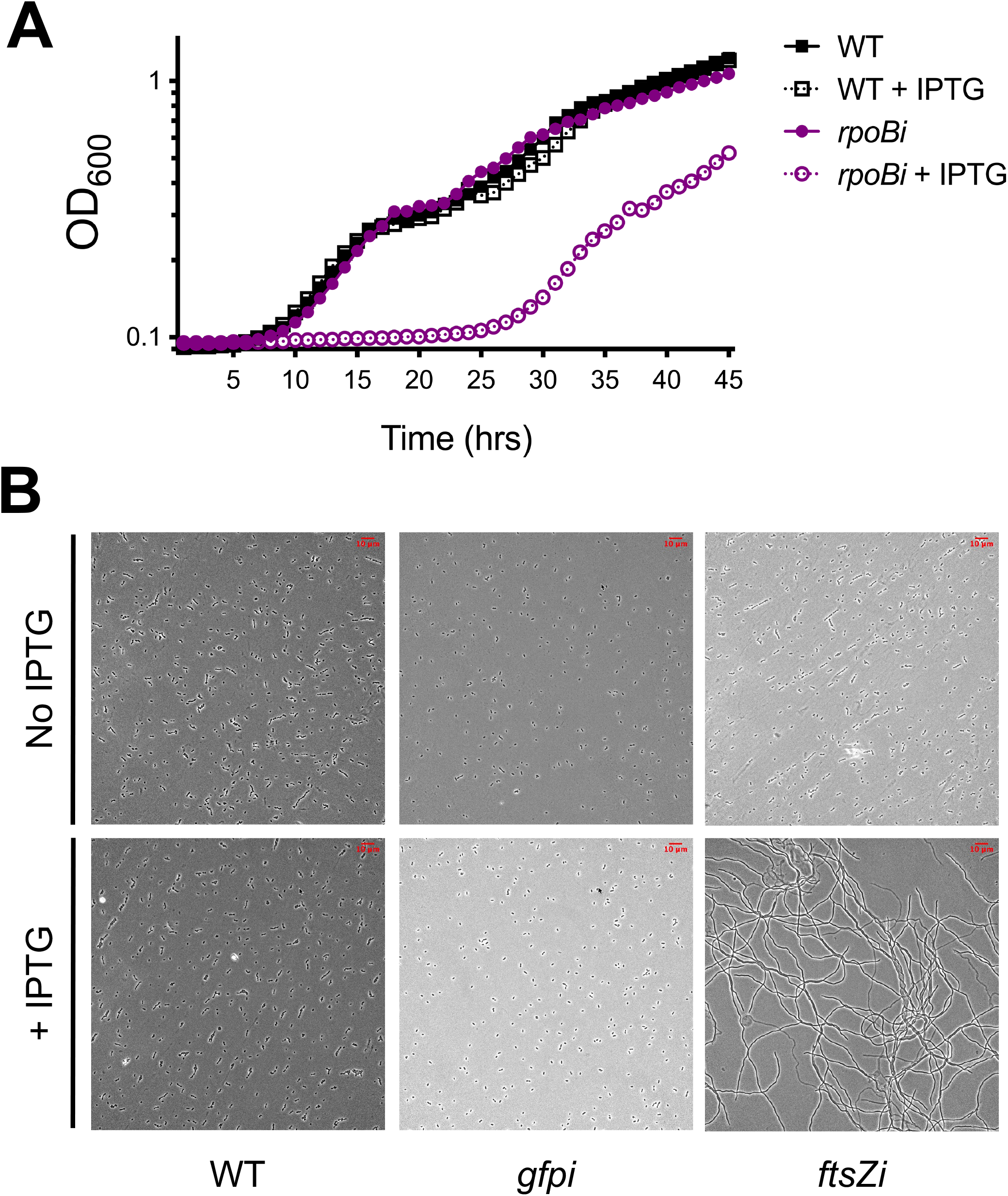
Targeting essential genes *rpoB* and *ftsZ* with CRISPRi. (A) Growth curves (OD_600_ over time) are shown for wild-type and *rpoBi* strains of *V. campbellii* in the presence and absence of 100 μM IPTG. One data set of three biological replicates is shown. (B) Micrographs of wild-type, *gfpi*, and *ftsZi* strains at 8 hours post-induction. Scale bars (10 μm) are shown.

To further test the utility of the Mobile-CRISPRi system, we targeted the divisome gene, *ftsZ*, and observed the effect of knockdown on cell growth and shape [20, 21]. As expected, severe filamentation of the cells was observed throughout the entire population at 8 hours post-IPTG induction, consistent with a defect in division (Fig. 3B). The *gfpi* control strain lacked any observable difference from the wild-type cells. From these experiments, we conclude that Mobile-CRISPRi can be used to target essential genes to observe phenotypes in live cells.

### *Mobile-CRISPRi functions in multiple* Vibrio *species*

To test the efficacy of Mobile-CRISPRi in other *Vibrio* species, we designed and cloned sgRNAs that target the LuxR homolog in *V. cholerae, V. vulnificus, and V. parahaemolyticus*. All three of these species are not naturally bioluminescent, so we used an ectopic fluorescent reporter plasmid (P_lux_-*gfp*) that measures LuxR-dependent *luxCDABE* promoter activation (pCS42 – see Fig. 4A for simplified diagram). Repression of the genes encoding the *V. vulnificus* (SmcR) and *V. parahaemolyticus* (OpaR) LuxR homologs in their native backgrounds resulted in *gfp* repression similar to complete deletion of the same gene (Fig. 4A). We also attempted this assay in *V. cholerae*, however, after attempting to clone in three different guides targeting the LuxR homolog in *V. cholerae* (HapR), there was still no successful repression of *gfp* expression. To determine if this was due to *V. cholerae* being incompatible with the system, we tested a *gfp* sgRNA in all our *Vibrio* species. We observed that in all four species, fluorescence was significantly reduced upon induction of the CRISPRi suite (Fig. 4B). However, considering those data with our initial issues with repressing *hapR* expression, suggests that the Mobile-CRISPRi system is functional in *V. cholerae* but that additional optimization may be required.

**Fig. 4.**
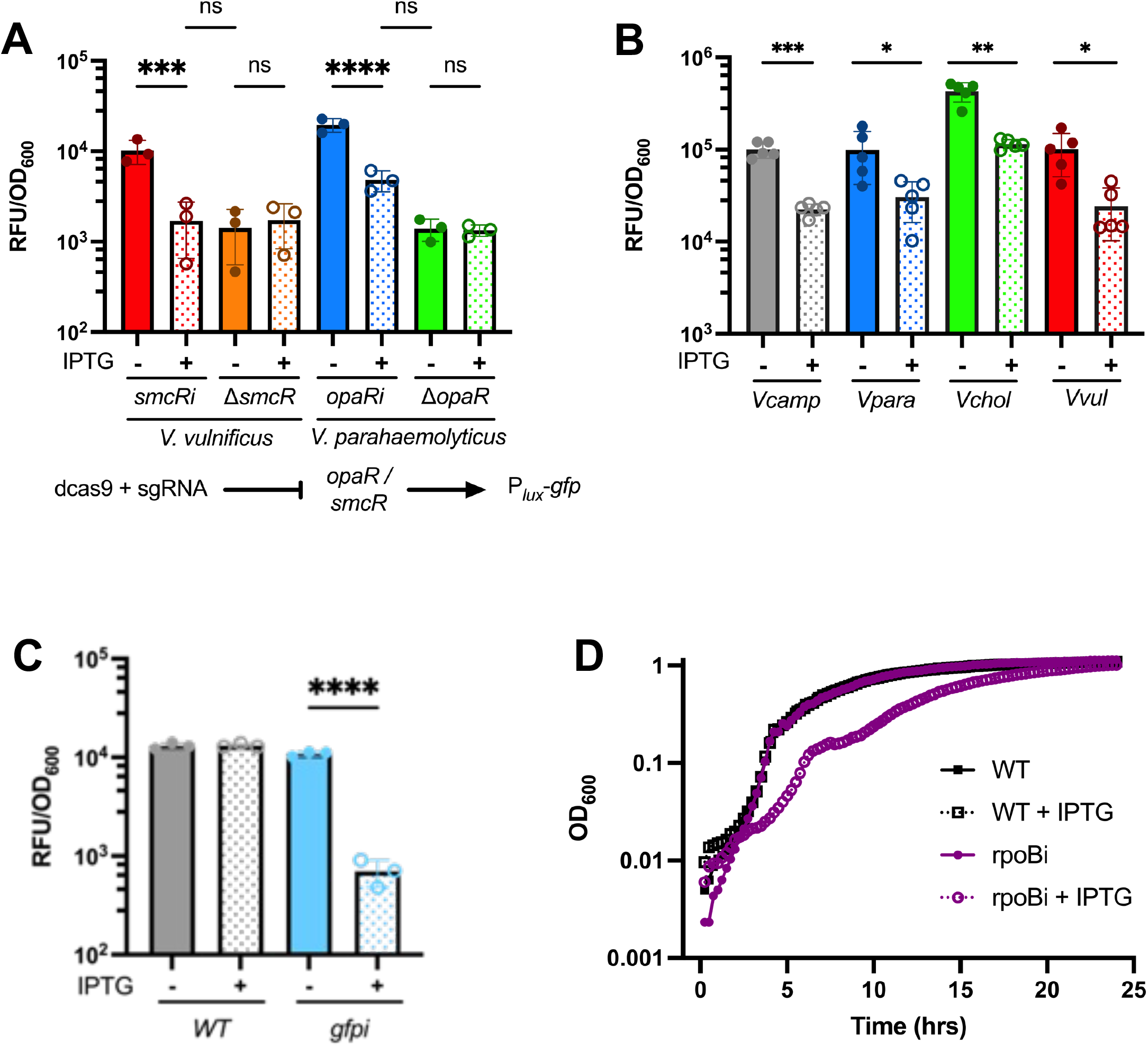
Mobile CRISPRi functions in multiple *Vibrio* species. (A) Fluorescence endpoint production (RFU/OD_600_) at 15 hours is shown for strains of *Vibrio* in wild-type and *luxR* deletion backgrounds containing a CRISPRi plasmid with guide RNA targeting the *luxR* homolog in each species in the presence (+) or absence (-) of 100 μM IPTG. Below the graph is a simplified diagram depicting the assay set up. (B) Fluorescence endpoint production (RFU/OD_600_) at 15 hours is shown for strains of *Vibrio* in wild-type backgrounds containing a CRISPRi plasmid with guide RNA targeting *gfp* in each species in the presence (+) or absence (-) of 100 μM IPTG. Both (A) and (B) utilize a reporter with *gfp* transcriptionally fused to a LuxR activated locus (P_*luxCDABE*_). (C) Fluorescence endpoint production at 15 hours is shown for *V. fischeri* ES114 in Mobile-CRISPRi backgrounds with (*gfpi*) or without (WT) sgRNA targeting *gfp* in the presence (+) or absence (-) of 100 μM IPTG. (D) Growth curves (OD_600_ over time) are shown for wild-type and *rpoBi* strains of *V. fischeri* ES114 in the presence and absence of 100 μM IPTG. One data set of three biological replicates is shown. Statistical analysis: normally distributed data (D’Agostino and Pearson test) were analyzed by two-way analysis of variance (*n*=3; Tukey’s multiple comparisons test; *, *p*=0.05; **, *p*=0.01; ***, *p*=0.001; ****, *p*=0.0001; ns=not significant). Error bars represent Standard Deviation.

To test additional targets in another *Vibrio* species, we focused on *V. fischeri*. We targeted *rpoB* in a fashion like that conducted for *V. campbellii*, as *rpoB* is also predicted to be an essential gene in *V. fischeri* ES114 [21]. Upon induction of the *V. fischeri rpoBi* strain with IPTG, we observed a significant growth lag, which we did not observe upon addition of IPTG in a control wild-type strain (Fig. 4D). We also note that the uninduced *rpoBi* strain does not exhibit any growth lag in this experiment nor in the analogous *V. campbellii* experiment (Fig. 3A, 4D), suggesting that the system is not leaky enough to convey a fitness deficit in this context, making it especially suitable to probe functions of essential genes. Additionally, targeting *gfp* was tested in *V*. fischeri. We observed that Mobile-CRISPRi was, once again, able to significantly reduce the expression of GFP (Fig. 4C). Overall, these experiments demonstrate that Mobile-CRISPRi is effective in *V. campbellii, V. cholerae, V. vulnificus, V. fischeri, and V. parahaemolyticus*.

## Discussion

This work focused on developing the Mobile-CRISPRi suite for use across *Vibrio* bacteria with the objective of determining the specificity, efficacy, and versatility of this system for studying *Vibrio* biology. We targeted multiple non-essential and essential genes to test the parameters of this assay across five *Vibrio* species. Our data demonstrated that Mobile-CRISPRi can deplete gene expression such that the resulting phenotypes mimic deletion of the same gene. Additionally, we show that the Mobile-CRISPRi suite does not impact the growth of *V. campbellii* when targeting a locus expected to be neutral to growth (Fig. 1A, Fig. S2). Consistent with the literature in other systems, the greatest and most consistent repression is achieved by targeting the 5’ end of the target’s open reading frame (Fig. 2) [6, 23, 25]. Through targeting of both *rpoB* and *ftsZ*, we show that Mobile-CRISPRi can deplete expression of essential genes and identify their physiological impacts on the cell (Fig. 3A, 3B, 4D). The *rpoBi* results in *V. fischeri* parallel those obtained in *V. campbellii*, including the eventual recovery in growth (Fig. 3A, 4D). These data in combination with the *gfpi* experiments (Fig. 4) point to significant consistency in the use of CRISPRi across the *Vibrionaceae*. Furthermore, the *V. fischeri* work in the study was conducted in a different laboratory from that of the other species, and the congruity in the results obtained suggests that adoption of the technology by new groups will yield reproducible results.

Mobile-CRISPRi readily functions in a variety of *Vibrio* species (Fig. 4), demonstrating its potential utility in studying non-model organisms that typically lack robust genetic tool kits. The utility of CRISPRi in *Vibrio* is further supported by a plasmid tool kit recently developed for *V. parahaemolyticus* [25]. However, the Mobile-CRISPRi previously published and utilized here expands this knockdown technique by allowing for stable chromosomal integration of the necessary machinery. This stable integration prevents cell-to-cell deviations due to plasmid copy number and allows for the system to be reliably maintained within a host extending its utility into infection and colonization assays [9]. Given the extensive use of *Vibrio* spp. in animal model systems to study human diseases, aquatic diseases, and animal symbioses, stable chromosomal integration presents a substantial benefit.

The development of genetic tools for model and non-model organisms is essential for elucidation of gene function. Transposon mutagenesis is one of the most powerful tools used in bacterial genetics. Similarly, we anticipate that the Mobile-CRISPRi suite will enable new and fundamental experiments in which both essential and non-essential genes can be probed in screens for function. Indeed, this system has already been proven to revolutionize large-scale genetic screens using oligonucleotide-based spacer libraries that target all essential [26] or all annotated genes in the genome [27]. Experiments in model systems such as *V. cholerae* and *V. fischeri* will expand the toolsets available to enable host-microbe studies of gene function *in vivo*.

## Methods

### Bacterial strains and plasmids

All oligos utilized for cloning in this study can be found in Table S1. All plasmids and bacterial strains used herein are listed in Tables S2 and S3. *V. campbellii* strains were grown in Luria Marine (LM) medium (Lysogeny broth with an additional 10 g per L of NaCl) at 30 °C, and *V. fischeri* strains were grown in Lysogeny broth salt medium (LBS; LM that is buffered with 50 mL 1 M pH 7.5 Tris per L) at 25 °C. All other strains were grown in Lysogeny broth (LB) at 37 °C. When required, the appropriate media was supplemented with kanamycin (100 µg/ml), gentamycin (100 µg/ml), ampicillin (100 µg/ml), chloramphenicol (10 µg/ml), or isopropyl β-D-1-thiogalactopyranoside (IPTG) at 100 µM unless otherwise noted. Plasmids were transferred from *E. coli* to *Vibrio* strains via conjugation on LB plates or LBS for *V. fischeri*. Exconjugants were selected for using polymyxin B (50 µg/ml) and any additional antibiotics required.

### sgRNA Design

The sgRNA targets were designed as described in Banta *et al*. [2].

To construct a comprehensive list of sgRNA sequences, a command-line interpreter was utilized. Anaconda Navigator (Conda) was downloaded onto a Unix compatible computer from https://docs.conda.io/projects/conda/en/latest/. A Conda environment was prepared through a script taken directly from Banta *et al*. Further information is available through Github documentation (https://github.com/ryandward/sgrna_design) at the sgrna_design repository and the Banta *et al*. protocol [2].

The genes of interest were then identified by their locus tag within the list. Target sgRNAs were selected from this list and ordered through IDT. An example oligo pair to produce a proper sgRNA can be found below:

Top Oligo: tagtAGAGGGGAAAGCCTAGTACG

Bottom Oligo: aaacCGTACTAGGCTTTCCCCTCT

Lower case indicates complementarity with a BsaI sticky end that is generated in pJMP1339. The upper case indicates sgRNA sequence. The Top Oligo refers to the sequence from the target list. The Bottom Oligo is the reverse complement of this sequence.

### Molecular cloning methods

The pJMP1339 plasmid DNA was digested using the BsaI HF-v2 Restriction endonuclease (NEB) for four hours at 37 ºC, purified using a Qiagen PCR Purification kit, and quantified using a BioTek Synergy H1 plate reader. The 5’-phosphate was added via T4 polynucleotide kinase (NEB) at 37 ºC for 30 minutes. The T4 kinase was then heat inactivated through incubation at 65 ºC for 20 minutes. The oligos were then annealed by incubating them at 95 ºC for five minutes and allowing them to cool to room temperature. This final solution was diluted 1:40. Ligations of the dsDNA oligos to the digested pJMP1339 were performed with T4 ligase (NEB) in a 10 µl reaction volume with 50 ng of digested pJMP1339 and 2 µl of diluted oligos, and the reaction incubated for 2 hours at room temperature followed by heat-inactivation for 20 minutes at 65 ºC. Reactions were transformed into electrocompetent *Escherichia coli* S17-λpir cells or *E. coli* WM6026 for *V. fischeri [2]*. Plasmids were sequenced to confirm (Eurofins).

### *Plasmid conjugation into* Vibrio

Mobile-CRISPRi machinery was conjugated into *V. fischeri* using a tri-parental mating of DAP auxotrophic donors for Tn7 transposition: sJMP3049 (*E. coli attTn7*::*acrIIA4, recA1, dapA*::*pir*) for cloning the Mobile-CRISPRi with a unique sgRNA-spacer, pTn7C1 as the transposase donor, and *V. fischeri* as the recipient. 100 µl of overnight cultures for each strain was added to 700 µl of LBS and centrifuged at 8,000 x g for 2 minutes. Then, cells were resuspended in 1 ml of LBS and centrifuged again at 8,000 x g for 2 minutes. Centrifuged cells were resuspended on 10 µl of LBS DAP and spotted on LBS-DAP agar plates. The plate was incubated lid up at 25 ºC overnight. Each sample was resuspended in 750 µl of LBS and plated on selective LBS-kanamycin agar plates and incubated at room temperature for 2 days. Eight colonies were tested via colony PCR. Three successful inserts were grown overnight for cryopreservation and downstream assays.

When attempting this protocol in some of the other *Vibrio* strains, the conjugative efficiency was insufficient for successful integration of the system. Therefore, we optimized the mating protocol for the other *Vibrio*.

Plasmid constructs were introduced into the other *Vibrio* cells via quadra-parental mating. For example, to mate a plasmid into *V. campbellii* BB120, the following strains were used: BB120 (*Vibrio* recipient), pJMP1039 (encoding the Tn7 transposase) [2, 3], pRK600 (mating helper strain) [22], and the S17-*1*λ*pir* strain containing the plasmid of interest (*e*.*g*., pJMP1339 encoding the Mobile-CRISPRi suite onto which the sgRNA had been cloned). 1 ml of each culture was centrifuged at 8,000 x g for 5 min and resuspended in non-selective LB media. 10 µl of each suspension was then dispensed in four total co-spots contained on an LB agar plate. The plate was then stored lid up at 30 ºC overnight. Each co-spot from a given plate was resuspended in the same 1 ml of LM media, and 100 µl of this suspension was streaked for isolation on an LM plate with 100 µg/ml kanamycin (selection for integration of Mobile-CRISPRi machinery) and 50 µg/ml polymyxin B (selection for Vibrio) and incubated overnight at 30 ºC. Three of the resulting colonies were then tested for successful Mobile-CRISPRi insertion via colony PCR. One of the successful colonies was inoculated overnight in the appropriate liquid culture and cryopreserved for downstream assays.

### Bioluminescence and GFP fluorescence measurements

The bioluminescence assays were performed in black-welled, clear bottom 96-well sterile plates. Selective media (240 µl) was added to each well, and 30 µl of 1 μM IPTG stock or dH_2_O was added (unless otherwise stated). Overnight cultures were diluted 1:1,000 in appropriate selective media, and 30 µl of these dilutions were added to the appropriate wells for a final volume of 300 µl. The plates were incubated and read using a BioTek Cytation3 plate reader and Gen5 Microplate reader software. The plate reader incubated the plate at 30 ºC with a continuous double orbital kinetic setting that took a reading every 30 minutes for 22 hours, recording both OD_600_ and bioluminescence (lux). Each experiment had three biological replicates, each with two technical replicates per sample. The data were analyzed by averaging the technical replicates for both OD_600_ and lux, and the data were plotted as relative lux units (lux/OD_600_).

For measuring GFP fluorescence in *Vibrio* species other than *V. fischeri*, overnight cultures grown in selective media were centrifuged for 5 minutes at 8,000 x g and resuspended in 2x PBS. After resuspension, 300 μl was added to black-welled, clear bottom 96-well sterile plates and measured using a BioTek Cytation3 plate reader. For measuring GFP fluorescence in *V. fischeri*, 200 μl of overnight cultures grown in selective media was added to black-welled, clear bottom 96-well sterile plates and measured using a BioTek Synergy Neo2 plate reader.

## Acknowledgments

The authors thank the van Kessel lab for comments on the manuscript and helpful discussions. Research reported in this publication was supported by: 1) the National Institute of General Medical Sciences (NIGMS) of the National Institutes of Health (NIH) under award numbers R35GM124698 to JVK, R35GM150487 to JMP, R35GM148385 to MJM, and T32GM135066 to NRM. The content is solely the responsibility of the authors and does not necessarily represent the official views of the National Institutes of Health; 2) the National Science Foundation Graduate Research Fellowship Program to LJG.

## Conflicts of interest

JMP has filed for patents related to Mobile-CRISPRi technology. All other authors declare that they have no conflicts of interest.

