## Supplemental material for "Mobile-CRISPRi as a powerful tool for modulating *Vibrio* gene expression"

**Table S1 Strains.**

| ***Strain*** | ***Species*** | ***Genotype*** | ***Description*** | ***References*** |
|---|---|---|---|---|
| LG011 | *V. campbellii* | pJMP1339 / WT | Empty Mobile-CRISPRi integrated into the genome | This study |
| LG020 | *V. campbellii* | pLG04 | rpoBi (18) | This study |
| LG022 | *V. campbellii* | pLG02 | luxRi (25) | This study |
| LG031 | *V. vulnificus* | pLG014 + pCS042 | smcRi (108) with plux::gfp reporter | This study |
| LG032 | *V. vulnificus* | Δ*smcR* pLG014 + pCS042 | smcRi (108) with plux::gfp reporter in a Δ*smcR* background | This study |
| LG035 | *V. parahaemalyticus* | pLG016 + pCS042 | opaRi (208) with plux::gfp reporter | This study |
| LG036 | *V. parahaemalyticus* | Δ*opaR* pLG016 + pCS042 | opaRi (208) with plux::gfp reporter in a Δ*opaR* background | This study |
| LG047 | *V. campbellii* | pMT05 | gfpi (137) | This study |
| LG048 | *V. campbellii* | pMT05 + pCS042 | gfpi (137) with plux::gfp reporter | This study |
| LG083 | *V. parahaemalyticus* | pMT05 + pCS042 | gfpi (137) with plux::gfp reporter | This study |
| LG084 | *V. parahaemalyticus* | Δ*opaR* pMT05 + pCS042 | gfpi (137) with plux::gfp reporter in a Δ*opaR* background | This study |
| LG085 | *V. cholerae* | pMT05 + pCS042 | gfpi (137) with plux::gfp reporter | This study |
| LG086 | *V. cholerae* | Δ*hapR* pMT05 + pCS042 | gfpi (137) with plux::gfp reporter in a Δ*hapR* background | This study |
| LG087 | *V. vulnificus* | pMT05 + pCS042 | gfpi (137) with plux::gfp reporter | This study |
| LG088 | *V. vulnificus* | Δ*smcR* pMT05 + pCS042 | gfpi (137) with plux::gfp reporter in a Δ*smcR* background | This study |
| MT03 | *V. campbellii* | pMT01 | luxCi (17) | This study |
| MT06 | *V. campbellii* | pMT02 | luxCi (1430) | This study |
| MT07 | *V. campbellii* | pMT03 | luxCi (950) | This study |
| MT09 | *V. campbellii* | pMT04 | ftsZi (13) | This study |
| MT16 | *V. campbellii* | pMT06 | luxRi (108) | This study |
| MT17 | *V. campbellii* | pMT07 | luxRi (318) | This study |
| MT18 | *V. campbellii* | pMT08 | luxRi (478) | This study |
| MT19 | *V. campbellii* | pMT09 | luxRi (606) | This study |
| sJMP3049 | *E. coli* | WM6026 | | [1] |
| sJMP2846 | *E. coli* | WM6026/pJMP2846 | | [2] |
| sNRM040 | *E. coli* | WM6026 / pNRM40 | | This study |
| sNRM046 | *V. fischeri* | ES114 (MJM1100) / pJMP2846 | Empty Mobile-CRISPRi integrated into the genome | This study |
| sNRM047 | *V. fischeri* | ES114 (MJM1100) / pNRM40 | rpoBi (34) | This study |
| sNRM044 | *V. fischeri* | ES114 (MJM1100) / pJMP2822 | Empty Mobile-CRISPRi integrated into the genome. Expressing GFP. | This study |

| | | | | |
|---|---|---|---|---|
| sNRM045 | *V. fischeri* | ES114 (MJM1100) / pJMP2834 | gfpi expressing GFP. | This study |

**TS1.** Strains with Mobile-CRISPRi systems are denoted as (gene of interest) followed by an i and the open reading frame targeting position in parentheses.

**Table S2 Plasmids.**

| ***Plasmid*** | ***Genotype*** | ***Description*** | ***Reference*** |
|---|---|---|---|
| pLG001 | luxCi (272) | Mobile-CRISPRi for *luxC* interference. ORF position 272 | This study |
| pLG002 | luxRi (25) | Mobile-CRISPRi for *luxR* interference. ORF position 25 | This study |
| pLG008 | rpoBi (18) | Mobile-CRISPRi for *rpoB* interference. ORF position 18 | This study |
| pLG014 | smcRi (108) | Mobile-CRISPRi for *smcR* interference. ORF position 108 | This study |
| pLG016 | opaRi (208) | Mobile-CRISPRi for *opaR* interference. ORF position 208 | This study |
| pJMP1339 | dCas9 | Mobile-CRISPRi machinery without a guide for cloning | [2] |
| pJMP1039 | Tn7 Transposase | Transposase for insertion of Mobile CRISPRi machinery | [2] |
| pRK600 | R Factor | Helper plasmid for conjugation of CRISPRi into target strain | [3] |
| pMT01 | luxCi(17) | Mobile-CRISPRi for *luxC* interference. ORF position 17 | This study |
| pMT02 | luxCi(1430) | Mobile-CRISPRi for *luxC* interference. ORF position 1430 | This study |
| pMT03 | luxCi(950) | Mobile-CRISPRi for *luxC* interference. ORF position 950 | This study |
| pMT04 | ftsZi(13) | Mobile-CRISPRi for *ftsZ* interference. ORF position 13 | This study |
| pMT05 | gfpi(137) | Mobile-CRISPRi for *gfp* interference. ORF position 137 | This study |
| pMT06 | luxRi(108) | Mobile-CRISPRi for *luxR* interference. ORF position 108 | This study |
| pMT07 | luxRi(318) | Mobile-CRISPRi for *luxR* interference. ORF position 318 | This study |
| pMT08 | luxRi(478) | Mobile-CRISPRi for *luxR* interference. ORF position 478 | This study |
| pMT09 | luxRi(606) | Mobile-CRISPRi for *luxR* interference. ORF position 606 | This study |
| pJMP2846 | dCas9 | Mobile-CRISPRi empty sgRNA vector | [2] |
| pNRM40 | rpoBi (34) | Mobile-CRISPRi for *rpoB* interference in *V. fischeri* ES114. ORF position 34 | This study |
| pJMP2822 | dCas9 | Mobile-CRISPRi expressing GFP empty sgRNA vector | [2] |
| pJMP2834 | gfpi | Mobile-CRISPRi for *gfp* interference | [2] |

**TS2.** pJMP2846 was used for cloning in the *Vibrio fischeri* and pJMP1339 was used for all of the other *Vibrio* evaluated.

### Table S3 Oligos.

| *Primer* | *Sequence* | *Description* |
|---|---|---|
| LJG016 | tagtGCTTCGAGTTTCGCCATTTG | luxCi (272) Top Oligo |
| LJG017 | aaacCAAATGGCGAAACTCGAAGC | luxCi (272) Bottom Oligo |
| LJG022 | gccttctcgtctggtcagtttcacctg | Sequencing Primer for sgRNA inserts |
| LJG068 | gcgatctactacaccatcccaatgc | CRISPRi insertion confrimation, forward |
| LJG070 | atggtggaaaatggccgcttttc | CRISPRi insertion confrimation, reverse |
| LJG107 | tagtATTGTGAATTTCCTTGTTTA | luxCi (1430) - top |
| LJG108 | aaacTAAACAAGGAAATTCACAAT | luxCi (1430) - bottom |
| LJG109 | tagtTCAAAATTTTGCTTTGATTT | luxCi (950) - top |
| LJG110 | aaacAAATCAAAGCAAAATTTTGA | luxCi (950) - bottom |
| LJG111 | tagtCTTACGGATGCGCTTTTTCT | rpoBi (18) - top |
| LJG112 | aaacAGAAAAAGCGCATCCGTAAG | rpoBi (18) - bottom |
| LJG195 | tagtGGCCACAAGTTCTCTGTCAG | gfpi (90) - top |
| LJG196 | aaacCTGACAGAGAACTTGTGGCC | gfpi (90) - bottom |
| LJG199 | tagtTTGCAAAGAGACCTAGAACT | opaRi (28) - top |
| LJG200 | aaacAGTTCTAGGTCTCTTTGCAA | opaRi (28) - bottom |
| LJG233 | tagtCAATATCCGCGTGACCACCA | luxRi (108) - top |
| LJG234 | aaacTGGTGGTCACGCGGATATTG | luxRi (108) - bottom |
| LJG235 | tagtAATCATTGCATTAGTGATGT | luxRi (318) - top |
| LJG236 | aaacACATCACTAATGCAATGATT | luxRi (318) - bottom |
| LJG237 | tagtAATAGATTCGCCAAGTGTTC | luxRi (478) - top |
| LJG238 | aaacGAACACTTGGCGAATCTATT | luxRi (478) - bottom |
| LJG239 | tagtTTATTTTTTAGTGATGTTCA | luxRi (606) - top |
| LJG240 | aaacTGAACATCACTAAAAAATAA | luxRi (606) - bottom |
| LJG259 | tagtAGCGGAGATAAGCGAGTTCG | smcRi (25) - top |
| LJG260 | aaacCGAACTCGCTTATCTCCGCT | smcRi (25) - bottom |
| MPT01 | tagtTCGTCAGACATTTCCATCAT | ftsZi (13) - top |
| MPT02 | aaacATGATGGAAATGTCTGACGA | ftsZi (13) - bottom |
| 818_VF_2414_T | tagtGACGCTTACCAAAATCCTTA | rpoBi - top |
| 819_VF_2414_B | aaacTAAGGATTTTGGTAAGCGTC | rpoBi - bottom |

**TS3.** All oligos listed above were generated as part of this study.

**Figure S1.**

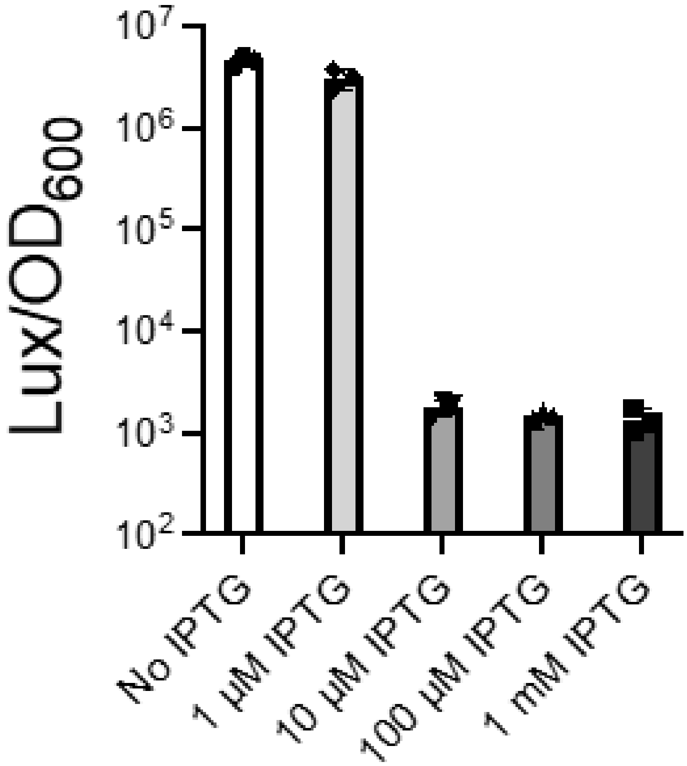


**Fig. S1.** IPTG titration with the *luxRi* strain of *V. campbellii* BB120. Error bars represent standard deviation (*n*=3).

**Figure S2.**

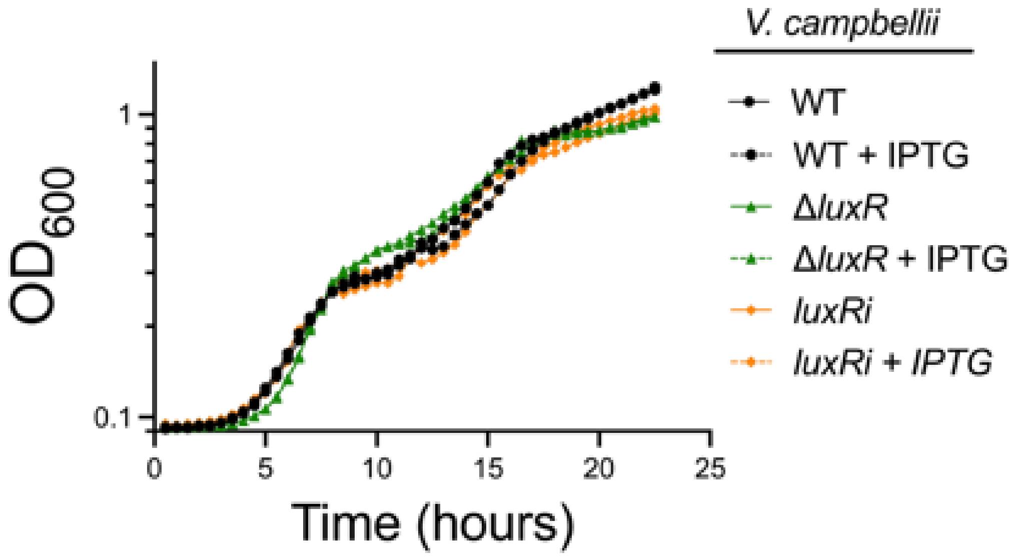


**Fig. S2.** Growth curves of *V. campbellii* BB120 strains. Representative of *n*=3.

**References.**


1. Ward, R.D., et al., *Essential gene knockdowns reveal genetic vulnerabilities and antibiotic sensitivities in Acinetobacter baumannii.* mBio, 2023.
2. Banta, A.B., et al., *Programmable Gene Knockdown in Diverse Bacteria Using Mobile-CRISPRi.* Curr Protoc Microbiol, 2020. **59**(1): p. e130.
3. Kessler, B., V.d. Lorenzo, and K. Timmis, *A general system to integrate lacZ fusions into the chromosomes of gram-negative eubacteria: regulation of the Pm promoter of the TOL plasmid studied with all controlling elements in monocopy.* Mol Gen Genet, 1992. **233**: p. 293-301.
